# Structure-guided stabilization improves the ability of the HIV-1 gp41 hydrophobic pocket to elicit neutralizing antibodies

**DOI:** 10.1101/2022.12.22.521671

**Authors:** Theodora U. J. Bruun, Shaogeng Tang, Graham Erwin, Lindsay Deis, Daniel Fernandez, Peter S. Kim

## Abstract

The hydrophobic pocket found in the N-heptad repeat (NHR) region of HIV-1 gp41 is a highly conserved epitope that is the target of various HIV-1 neutralizing monoclonal antibodies. Although the high conservation of the pocket makes it an attractive vaccine candidate, it has been challenging to elicit potent anti-NHR antibodies via immunization. Here, we solved a high-resolution structure of the NHR mimetic IQN17, and, consistent with previous ligand-bound gp41 pocket structures, we observed remarkable conformational plasticity of the pocket. The high malleability of this pocket led us to test whether we could improve the immunogenicity of the gp41 pocket by stabilizing its conformation. We show that the addition of five amino acids at the C-terminus of IQN17, to generate IQN22, introduces a stabilizing salt bridge at the base of the peptide that rigidifies the pocket. Mice immunized with IQN22 elicited higher avidity antibodies against the gp41 pocket and a more potent, albeit still weak, neutralizing response against HIV-1 compared to IQN17. Stabilized epitope-focused immunogens could serve as the basis for future HIV-1 fusion-inhibiting vaccines.

## Introduction

HIV-1 infection occurs through the fusion of the HIV-1 membrane with the host cell membrane, a process that is mediated by the viral envelope glycoprotein Env, consisting of gp120 and gp41 trimers.^1–4^ After gp120 binds to its host cellular receptors, membrane fusion proceeds through a transient structure known as the pre-hairpin intermediate (PHI) in which gp41 inserts a hydrophobic fusion peptide into the host cell and in the process exposes two regions of gp41: the N-heptad repeat (NHR) and the C-heptad repeat (CHR).^2,5^ The collapse of the NHR and CHR into a stable trimer-of-hairpins is thought to drive membrane fusion.^4,6–8^ The trimer-of-hairpins, also referred to as a six-helical bundle, contains an internal coiled-coil trimer formed by parallel NHR helices against which three CHR helices pack in an antiparallel manner.

The structure of the HIV-1 gp41 post-fusion bundle revealed a prominent hydrophobic pocket within the NHR consisting of seven discontinuous residues that are highly conserved (L568, V570, W571, K574, Q575, Q577, R579).^1^ Across the 8,075 HIV-1 sequences in the Los Alamos database, these seven pocket residues are between 97 to 99% identical (**Fig. S1**). The mRNA nucleotides that encode the pocket residues are found in stem V of the Rev response element (RRE) (**Fig. S2**). The high conservation of the pocket residues at the nucleotide level is likely because the proper folding of the RRE is critical to the HIV-1 lifecycle.^9^

The NHR is a validated therapeutic target in humans since it is the binding site of the peptide enfuvirtide, an FDA-approved HIV-1 fusion inhibitor.^10,11^ Within the NHR, the gp41 pocket is a common binding site for other fusion inhibitors as well as neutralizing antibodies,^12–19^ and its extremely high level of conservation makes it an attractive vaccine target. Additionally, NHR-targeting antibodies have low levels of somatic hypermutation^20^ suggesting that it might be possible to elicit antibodies in a vaccine context without multistage, multicomponent, germline-targeting immunization strategies that are being widely pursued in HIV-1 vaccine development.^21–25^ Since the NHR is only transiently exposed in the PHI during the fusion process, eliciting an immune response against this target requires engineering a stabilized mimetic.^12,26–28^ One such PHI mimetic is the IQN17 peptide, which consists of a trimeric coiled-coil domain, GCN4-pIQI,^29^ fused to 17 NHR residues that include the gp41 pocket.^12^ However, vaccination of animals with NHR-presenting peptides has failed to elicit potently neutralizing antisera.^28,30–32^ The neutralizing activity of isolated anti-NHR monoclonal antibodies (mAbs) is also mainly limited to tier 1 viruses, which are by definition particularly sensitive to neutralization.^13,14,17,30,33^

Two recent discoveries suggest that the PHI warrants renewed investigation as a vaccine target. First, the activity of the pocket-targeting D5 antibody can be potentiated >1,000-fold by expression of FcγRI (CD64) on target cells in *in vitro* neutralization assays, leading to neutralization of tier 2 and tier 3 viruses, which are more representative of circulating strains.^34,35^ Since FcγRI is expressed on cells such as macrophages and dendritic cells that are prevalent in mucosal surfaces during the early stages of HIV-1 infection, it has been suggested that this potentiation might be relevant during sexual HIV-1 transmission.^31^ Second, an engineered version of D5 bearing additional amino acid substitutions in the heavy chain CDR loops, D5_AR, was able to neutralize tier 2 and tier 3 viruses.^20,35^ Although D5_AR is less potent than other broadly neutralizing antibodies, these results serve as an important proof of concept that antibodies targeting the PHI can broadly neutralize HIV-1 viruses.

Ligand-bound structures of the gp41 pocket, and the high-resolution apo structure of trimeric IQN17, indicate that the pocket is highly malleable.^1,12,14^ In particular, residue R579, which forms the base of the pocket as the last surface-exposed residue in IQN17, displays high conformational plasticity. We hypothesized that to elicit neutralizing antibodies, a rigidified pocket epitope would act as a better immunogen compared to a flexible pocket. A 2.0 Å X-ray crystal structure of IQN22, a C-terminally extended version of IQN17 containing five additional amino acids from the contiguous gp41 sequence, showed that the formation of an inter-helical salt bridge at the C-terminus stabilized the base of the pocket across all three faces of the trimer. Immunization with IQN22 in mice elicited antibodies with higher avidity to the gp41 pocket compared to antibodies from mice immunized with IQN17. Serum from mice immunized with the rigidified IQN22 peptide had a more potent, although still weak, neutralizing activity against HIV-1, compared to mice immunized with IQN17. In cells expressing FcγRI, antisera against stabilized-pocket peptides also weakly neutralized tier 1, tier 2, and tier 3 pseudoviruses. These results demonstrate that structure-guided design and rigidification of malleable epitopes can improve their immunogenicity. The stabilized peptides described here could guide future NHR-targeting HIV-1 vaccine efforts.

## Results

### The pocket of gp41 is highly malleable in the apo crystal structure

Initially, we sought to obtain a high-resolution structure of the apo gp41 pocket. Since the pocket of gp41 is only accessible in the transient pre-hairpin intermediate, an engineered stable mimetic is required for its proper display. For this, we used the previously described peptide, IQN17, which contains a soluble, trimeric coiled-coil fused to 17 amino acids of the NHR that include the gp41 hydrophobic pocket (residues 565-581 of HIV-1_HXB2_).^12^ IQN17 readily crystallized in the absence of a ligand and we were able to determine the X-ray crystal structure of the peptide to 1.2 Å (**Fig. 1a, Table S1**). The crystal contained one IQN17 trimer per asymmetric unit, with space group *I*222. The seven residues that form the pocket are V570, K574, and Q577 from one protomer, and L568, W571, Q575, and R579 from a neighboring protomer. Two of the three faces of the trimeric structure have crystal contacts that may influence the conformation of the pocket residue side chains. However, the pocket formed by chains A and C does not have crystal contacts, thus resembling a true ‘apo’ pocket. Strikingly, across all three faces, the residues are in different conformations from each other and from analogous sidechains in complex with a neutralizing antibody D5 (PDB ID 2CMR),^14^ a *D*-peptide inhibitor (PDB ID 1CZQ),^12^ and a CHR peptide derived from the gp41 ectodomain (PDB ID 1AIK) (**Fig. S3**).^1^ The average RMSD for these structures in superposition is 0.32 Å for the C_⍺_/C_β_atoms, indicating that changes in the pocket are predominantly attributable to different sidechain conformations. In particular, W571 and R579 show remarkable diversity in the adopted sidechain rotamers (**Fig. 1a,b**). Even though IQN17 ends at residue L581, the alpha-helical structure of the IQN17 chain places R579 at the C-terminus of the solvent exposed structure and at the base of the gp41 pocket. The B-factors of the backbone, C_⍺_, and side chain atoms of R579 are higher than average across the entire IQN17 structure, indicating conformational flexibility at this residue (**Table S2**).

**Figure 1.**
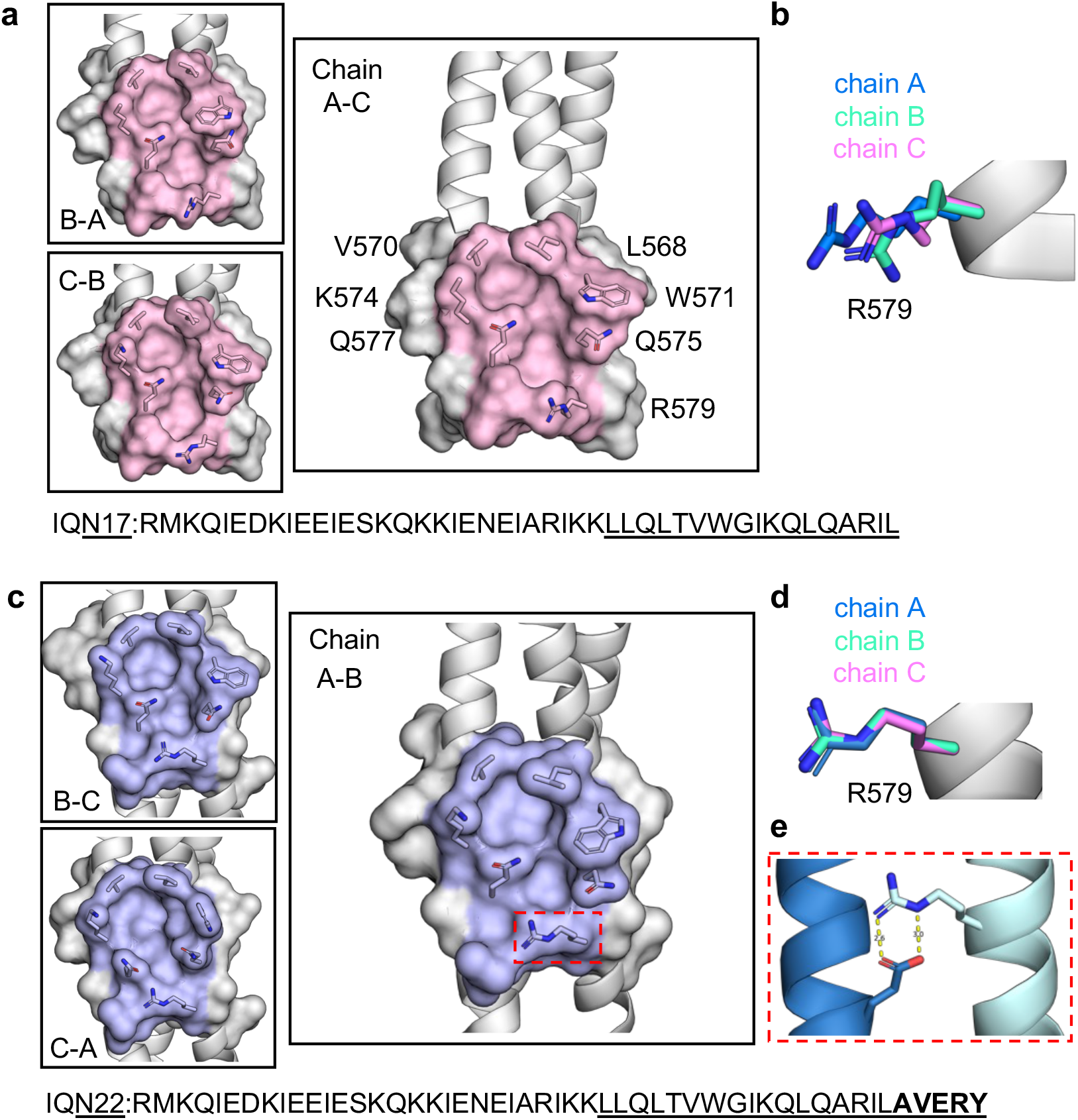
The gp41 pocket in the NHR mimetic IQN22 is rigidified compared to IQN17. **a**, A close-up view of the space-filling diagram of the gp41 pocket of IQN17 (pink) is shown with the seven conserved pocket residues labeled and the sidechains overlayed as sticks. **b**, Overlay of the side chain rotamers of R579 in each of the three chains of IQN17. **c**, A close-up view of the space-filling diagram of the gp41 pocket of IQN22 (lilac) with the pocket residue sidechains overlayed as sticks. **d**, Overlay of the side chain rotamers of R579 in each of the three chains of IQN22. **e**, R579 forms an inter-helical salt bridge with E584 on a neighboring chain.

### The presence of a salt bridge in the C-terminally elongated NHR mimetic IQN22 rigidifies the pocket

The crystal structure of IQN17 indicated that residue R579 displayed high conformational plasticity across the three faces of the trimeric peptide. Given the position of R579 near the C-terminus of IQN17, we hypothesized that further extending the C-terminus would stabilize the gp41 pocket epitope and thus improve the immunogenicity of the pocket. Using solid-phase peptide synthesis, we produced peptide IQN22 which contains the same residues as IQN17 and an additional five amino acids from the sequence of HIV-1_HXB2_ (residues 565-586) (**Table S3**). To interrogate the conformational changes caused by this extension, we solved a 2.0 Å X-ray crystal structure of IQN22 (**Fig. 1c, Table S1**). Similar to IQN17, the crystal contained one IQN22 trimer per asymmetric unit with space group *P*2_1_2_1_2_1_, and contained two crystal-contacted faces and one open pocket formed by chains A and B. Although W571 displays large side-chain flexibility, as seen in IQN17, R579 retains the same conformation across all three pockets of IQN22 (**Fig. 1d**). Additionally, B-factors for the R579 backbone, C_⍺_, and side chain atoms fall within the average values for the entire IQN22 structure (**Table S2**). Rigidification of R579 can be explained by the formation of an inter-helical salt bridge between R579 and the glutamic acid (E584) of the neighboring pocket-forming chain (**Fig. 1e**) that is not present in IQN17. Salt bridges are known to provide conformational specificity in proteins and to contribute to the stabilization of coiled-coil tertiary structures.^36,37^ The salt bridge between R579 and E584 has also been observed in the HIV-1 post-fusion six-helical bundle (PDB ID 1ENV).^3^

The structure of the D5-bound gp41 pocket reveals that R579 is engaged in a conformation similar to the position that R579 adopts in IQN22 (**Fig. S3**). In the structure of the pocket bound to the *D*-peptide inhibitor PIE12, R579 does not interact with any PIE12 residues and points away from the hydrophobic cavity of the pocket and towards the C-terminus of the NHR (**Fig. S3**). To gain insight into the in-solution conformation of IQN22, we measured binding of PIE12 or D5 Fab to IQN22 and IQN17 using bio-layer interferometry. We found that D5 Fab binds IQN17 and IQN22 with a *K_D_* of 14 and 5 nM, respectively, constituting an approximately threefold difference in affinity (**Table 1**). D5 Fab also had an increased association constant (k_on_) for binding to IQN22. The *K*_D_ for PIE12 binding to IQN17 and IQN22 was similar (3 nM v 2 nM), in line with the observation that R579 is important for binding of D5 but not implicated in the binding of PIE12 to the pocket (**Table 1, Fig. S3**). The increased affinity and association rate of D5 Fab binding to IQN22, as well as the structural evidence, suggests that the in-solution conformation of the IQN22 pocket is more rigid than that of IQN17.

**Table 1.** Summary of binding affinity and kinetic parameters for the binding of D5 Fab or PIE12 to IQN17 and IQN22.

|  | IQN17 |  |  | IQN22 |  |  |
| --- | --- | --- | --- | --- | --- | --- |
| | $K_D$<br>(nM) | $k_{on}$<br>($10^5 \text{ M}^{-1}\text{s}^{-1}$) | $k_{off}$<br>($10^{-3} \text{ s}^{-1}$) | $K_D$<br>(nM) | $k_{on}$<br>($10^5 \text{ M}^{-1}\text{s}^{-1}$) | $k_{off}$<br>($10^{-3} \text{ s}^{-1}$) |
| D5 Fab | $14 \pm 1.7$ | $5.7 \pm 0.35$ | $8.0 \pm 0.87$ | $5 \pm 1.9$ | $8.1 \pm 1.4$ | $4.1 \pm 2.2$ |
| PIE12 | $3.3 \pm 0.69$ | $1.9 \pm 0.56$ | $0.67 \pm 0.31$ | $2.4 \pm 0.96$ | $6.4 \pm 2.7$ | $1.5 \pm 0.56$ |

### Immunization with stabilized NHR peptides elicits higher avidity pocket-binding antibodies

Having shown that the pocket of IQN22 was stabilized compared to IQN17, we next explored our hypothesis that a rigidified immunogen is better at eliciting protective antibodies than a flexible one in a vaccine context. In addition to IQN17 and IQN22, we included another immunogen, IQN22AA, which replaced the five-amino acid (AVERY) extension in IQN22 with AVEAA (**Table S3**). This design was intended to minimize antibody responses to the immunodominant but non-neutralizing AVERY epitope^38^ while retaining the salt bridge and stabilization benefit from E584. The three immunogens, IQN17, IQN22, and IQN22AA were synthesized with an N-terminal CCGG motif (**Table S3**) and allowed to oxidize at neutral pH to form three interchain disulfide bridges, as previously described.^28,39^ We purified the resulting covalent trimers by high-performance liquid chromatography and evaluated their helicity by circular dichroism spectroscopy. All three immunogens showed high helicity, calculated as 100 %, 89 %, and 97 % helicity for IQN17, IQN22, and IQN22AA, respectively (**Fig. S4**).

Immunizations were performed in female Balb/c mice with five mice per group (**Fig. 2a**). Each mouse was immunized with 15 μg of peptide adjuvanted with alum (Allhydrogel, 500 μg) and CpG oligonucleotide (20 μg) on day 0, 21 and 42. Antisera were collected and analyzed post prime (day 21), post boost 1 (day 35), and post boost 2 (day 63).

**Figure 2.**
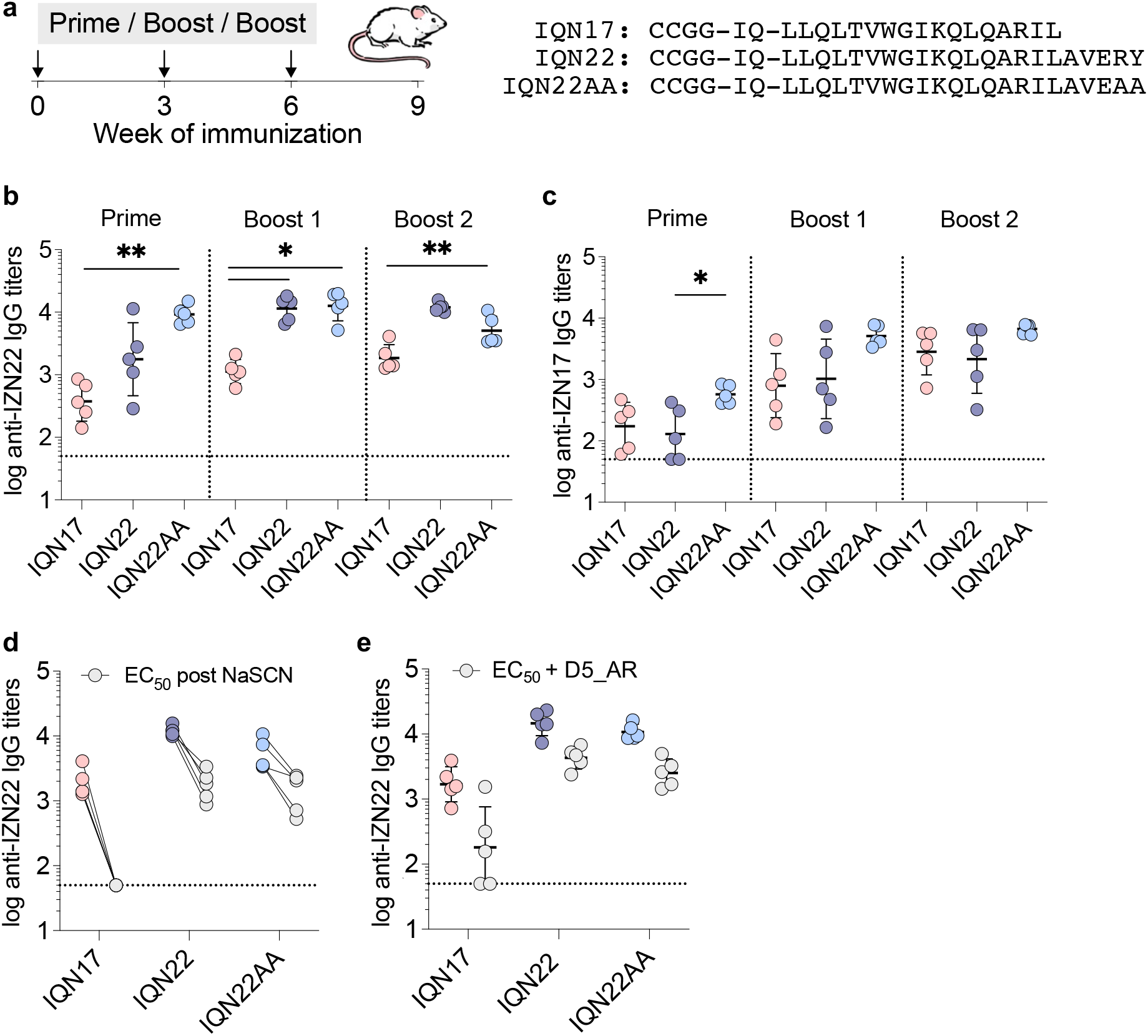
Immunization with stabilized-pocket immunogens elicits higher avidity antibodies against the gp41 pocket. **a**, Schematic of the three-dose immunization (n = 5 mice per group). Serum IgG titers to IZN22 (**b**) or IZN17 (**c**) at three time points. **d**, Serum binding titers to IZN22 at day 63 pre- and post- treatment with sodium thiocyanate (NaSCN). **e**, Serum binding titers to IZN22 at day 63 compared to serum binding titers in competition with pocket-binding antibody D5_AR. Data are presented as geometric mean ± s.d. of the log-transformed values. ELISAs were performed in biological duplicate. Statistical comparisons (**b, c**) were performed using Kruskal-Wallis ANOVA followed by Dunn’s multiple comparisons and only statistically significant p values are plotted. * = p ≤ 0.05, ** = p ≤ 0.01. The horizontal dotted line represents the limit of quantitation.

After a single dose, all mice developed antibody responses to the IQ domain as revealed by ELISA (**Fig. S5**). However, IQ-specific antibodies do not cross-react with IZ, another helical coiled-coil trimerization domain (**Fig. S5**).^26,40^ Thus, to specifically assess antibody titers to the pocket itself, we analyzed binding to IZN17 and IZN22 (**Table S3**), the analogous peptides of IQN17 and IQN22. While all groups developed antibody responses to the pocket after a single dose, which increased following each boost (**Fig. 2b,c**), the IQN22- and IQN22AA-immunized mice had significantly higher IgG titers to IZN22 than IQN17-immunized mice did (**Fig. 2b**). Titers against IZN17 were similar to those measured against IZN22, although the consistency within each group varied more widely (**Fig. 2c**). Antisera from mice immunized with IQN22AA displayed the most consistent antibody response against IZN17. We also assessed whether any antibodies were raised against AVERY and we observed no appreciable binding to the AVERY motif in mice immunized with IQN22 or IQN22AA (**Fig. S6**).

In addition to IgG titers, we assessed antibody avidity to the gp41 pocket by serum ELISAs in the presence of the chaotrope sodium thiocyanate (NaSCN).^41,42^ NaSCN displaces weakly bound antibodies so it allows a quantitative measure of the average strength of antibodies in a mixture. While NaSCN was able to displace the antibodies generated in IQN17-immunized mice, antibodies raised against the rigidified-pocket peptides remained bound at a greater proportion (**Fig. 2d**). Additionally, mice immunized with IQN22 or IQN22AA were better able to out-compete pocket binding antibody D5_AR in a competition ELISA (**Fig. 2e**). These results confirm our hypothesis that stabilization of the pocket epitope improves the immunogenicity of the NHR-displaying peptides.

### Antibodies elicited against the stabilized gp41 pocket neutralize HIV-1 in a pseudotyped lentiviral assay

To assess whether the antibodies raised against our peptide immunogens were functional, we used the well-established HIV-1 Env-pseudotyped lentiviral neutralization assay.^43^ Given that NHR peptides are known to induce weakly neutralizing antibodies,^28^ we used two strategies to enable detection of neutralizing responses. First, we performed neutralization assays using TZM-bl cells expressing FcγRI (TZM-bl/FcγRI) which have recently been shown to potentiate the neutralizing activity of PHI-targeting antibodies.^20,34^ Second, we used a tier 1 HIV-1 HXB2 virus that is particularly sensitive to PHI-targeting inhibitors due to the presence of an L565Q amino acid change.^44^

Neutralization of HXB2_L565Q_ at each of the three time points was strikingly different among groups (**Fig. 3a, Fig. S7a**). After the prime, we observed almost no neutralization with IQN17 and IQN22, whereas mice immunized with IQN22AA showed weak but significantly higher neutralization titers (NT_50_). Boosting increased the NT_50_ of IQN22- and IQN22AA-immunized mice while the neutralizing activity of IQN17-immunized mice barely increased above the baseline. After two boosts, mice immunized with IQN22AA had significantly higher NT_50_ than those immunized with IQN17 (**Fig. 3a**). Although not statistically significant, immunization with IQN22AA appeared to produce higher NT50 and a more consistent neutralization response compared to IQN22.

**Figure 3.**
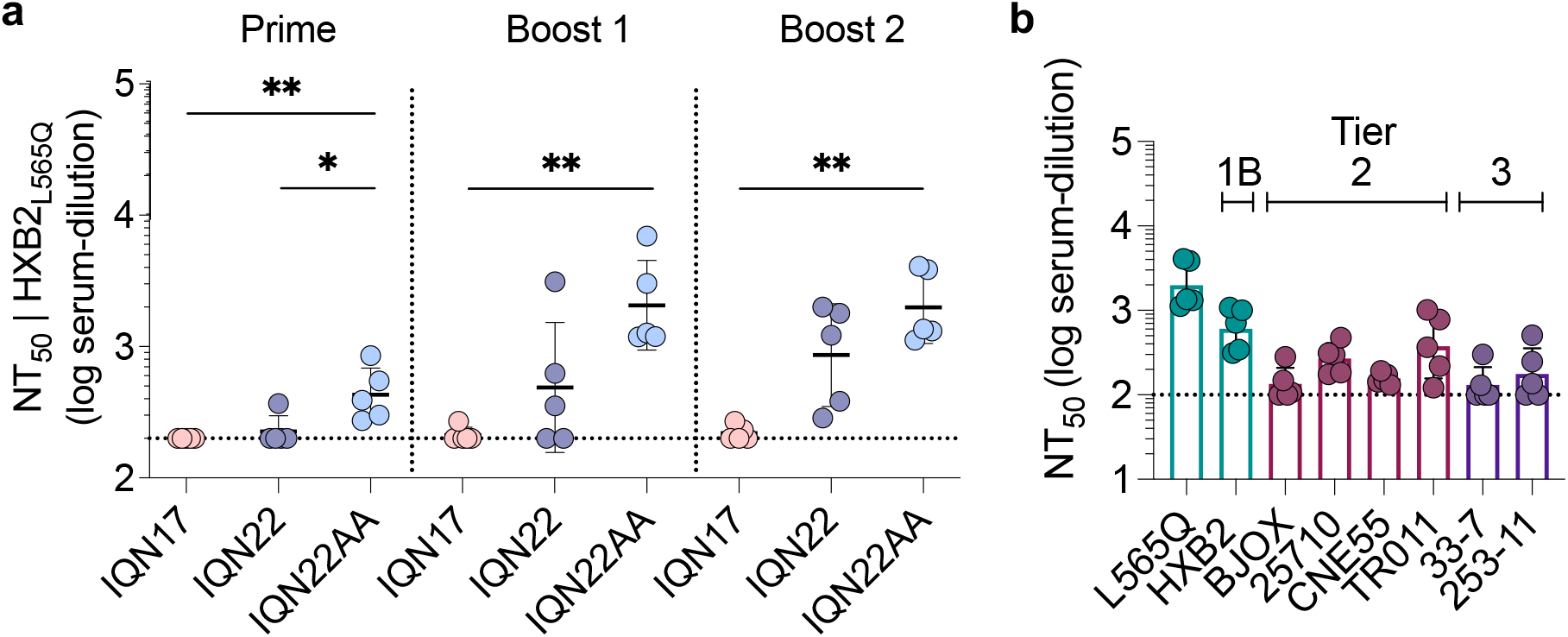
Immunization with stabilized gp41 pocket peptides elicits better neutralizing antibodies in an HIV-1 pseudotyped lentiviral assay. **a,**Serum neutralization titers (NT_50_) against HIV-1 HXB2_L565Q_ pseudotyped lentivirus over time. **b,** Serum NT_50_ from mice immunized with IQN22AA (post boost 2) against a panel of HIV-1 pseudotyped lentiviruses. Data are presented as geometric mean ± s.d. of the log-transformed values. Neutralization assays were performed in TZM-bl cells expressing FcγRI in biological duplicate. Comparison between the three groups at each time point was performed using a Kruskal Wallis ANOVA followed by a Dunn’s multiple comparisons test. P values less than 0.05 are considered significant and plotted. * = p ≤ 0.05, ** = p ≤ 0.01. The horizontal dotted line represents the limit of quantitation.

Finally, since the IQN22AA immunized mice had the highest and most consistent NT50 against HXB2_L565Q_, we assessed the neutralization breadth of this group. Antisera raised against IQN22AA weakly neutralized tier 2 and tier 3 HIV-1 viruses in TZM-bl/FcγRI cells (**Fig. 3b, Fig. S7b-d**), in line with the previous observation that PHI-targeting antibodies have weak but broad neutralization.^20,35^

## Discussion

Eliciting effective humoral immunity against HIV-1 continues to be a formidable challenge. The pre-hairpin intermediate has been the target of drug and vaccine efforts ever since its existence was first proposed decades ago.^45,46^ In particular, the pocket of gp41 is an attractive vaccine target because it is highly conserved across HIV-1 strains and because it is possible to raise neutralizing antibodies against this target^28,30–32,47^ without the use of germline-targeting strategies that employ multistage, multicomponent, immunization schemes.^21–25^

The high-resolution crystal structure of the apo pocket of IQN17, reported here, revealed remarkable flexibility of the side chains that differed across the three faces of the trimer and from analogous side chains in ligand-bound structures. This observation led us to hypothesize that a rigid pocket may act as a better immunogen compared to a flexible one. The structure of an elongated version of IQN17, IQN22, showed that the formation of a salt bridge between R579 and E584 rigidified the base of the pocket. The stabilized pocket of IQN22 more closely resembled the conformation of the D5-bound gp41 pocket. The higher affinity and *k_on_* for D5 Fab binding to IQN22 as compared to IQN17 suggests pre-stabilization of the D5-bound conformation.

Mouse immunization studies showed pocket-stabilized immunogens IQN22 and IQN22AA elicited antibodies that had higher avidity to the pocket and were better able to outcompete D5_AR for pocket binding. Antisera from mice immunized with stabilized peptides also had higher, albeit still weak, HIV-1 neutralization potencies compared to mice immunized with IQN17. With the expression of FcγRI on target cells we observe weak neutralization of HIV-1 viruses across all three tiers. This would especially be noteworthy if FcγRI plays a role in enhancing protection of cells that are infected in the mucosa and transmit virus to CD4+ T cells in the early stages of HIV-1 sexual transmission.^34,48–51^

Previous attempts to elicit neutralizing polyclonal antisera targeting the NHR showed that neutralization potency correlated with the concentration of D5-like antibodies.^28^ In line with this observation, our competition ELISA data with D5_AR suggest that the stabilized-pocket immunogens were better able to elicit pocket-binding D5-like antibodies. Although statistically non-significant, the neutralizing response was higher in mice immunized with IQN22AA compared to IQN22. Given that alanine has high helical propensity,^52,53^ it is possible that the addition of two alanine residues at the C-terminus of IQN22AA enhanced the helix-forming tendency of the immunogen, thus further stabilizing the C-terminus of the peptide.

Using a minimal NHR mimetic as an immunogen is an example of epitope-focusing,^54^ whereby antigens are engineered to promote the elicitation of antibody responses to a specific target of interest. However, peptides have short half lives *in vivo* and are generally weak immunogens.^55,56^ To increase the neutralizing activity over what we observed in this study, one could consider conjugating the peptides to carrier proteins or multimerizing on nanoparticles.^57–59^ Harnessing other vaccine advances such as hydrogel-based sustained release strategies for continuous immune stimulation, or the use of alternative adjuvants like saponin/MPLA may also increase the potency of NHR targeting vaccines.^60,61^

The structural, binding and immunization results presented here indicate that epitope conformation and rigidity can be engineered to bias the population of elicited antibodies in a vaccine context. Our findings also suggest that protein engineering can improve the gp41 pocket as a vaccine target and provide support that targeting the PHI is an orthogonal and complimentary approach to current HIV-1 germline-targeting vaccine efforts. More generally, the use of epitope-focusing and conformational rigidification to bias the immune response towards a particular subset of antibodies can be applied to the development of vaccines for other challenging infectious diseases.

## Methods

### Peptide synthesis and purification

Peptides were synthesized on a CSBio instrument using standard Fmoc-based solid-phase peptide synthesis. For each peptide, 250 μmol Rink amide AM resin (CSBio) or 250 μmol of NovaSyn TGR R resin (Sigma Aldrich) was used. NovaSyn TGR R resin has lower molecular weight loading and increased swelling properties and is preferrable for longer peptides. Coupling steps were performed with 1 mmol of amino acid (4X excess) for either 3 h at room temperature or 15 min at 60 °C in the presence of HBTU and *N,N*-diisopropylethylamine (DIEA). After synthesis, peptides were acetylated at the N-terminus manually using acetic anhydride. Peptides were cleaved from the resin using 95 % trifluoroacetic acid (TFA), 2.5 % water, 2.5 % triisopropylsilane (TIS) for 3-4 h at room temperature. For peptides containing cysteines the cleavage cocktail consisted of 94 % TFA, 2.5 % water, 1 % TIS, and 2.5 % 1,2-ethanedithiol (EDT). Cleaved peptides were precipitated in cold diethyl ether and then dissolved in a 1:4 mixture of acetonitrile and water and purified via high-performance liquid chromatography (HPLC) on a C18 semi-prep column using a water-acetonitrile gradient in the presence of 0.1 % trifluoroacetic acid. Fractions were collected and pooled based on liquid chromatography-mass spectrometry (LC/MS) analysis and then lyophilized and stored at −20 °C.

### Bio-layer interferometry

Bio-layer interferometry was performed on an Octet RED96^®^ system (Pall FortéBio) in 96-well flat-bottom black plates (Greiner). Streptavidin biosensor tips (Pall FortéBio) were pre-equilibrated in Octet buffer (1X DPBS [Gibco] with 0.1 % BSA and 0.05 % Tween-20) for 10 min before loading with biotinylated peptide (200 nM) to a load threshold of 0.7 nm. Peptide-loaded sensors were then dipped into known concentrations of ligand (PIE12 or D5 Fab) for an association step and then returned to the baseline well for a dissociation step. Binding curves were fit using the Pall FortéBio software using a 1:1 binding model, with background subtraction using a peptide-loaded sensor tip submerged into sample wells containing buffer only. Reported *K*_D_, *k*_on_, and *k*_off_ values are averages from fitted curves across multiple concentrations from at least two different independent experiments.

### Circular dichroism

Lyophilized peptide samples were suspended in 0.25X PBS buffer and concentrations were determined using a Nanodrop 2000 (ThermoFisher). CD spectra were collected using a JASCO J-815 CD spectrometer sampling every 0.5 nM between 260 nM and 180 nM. Three accumulations were collected and averaged for each sample. A cuvette containing buffer alone was run under the same conditions and the signal was subtracted from the sample runs. The data for each sample was converted from units of ellipticity (millidegrees) into units of mean residue ellipticity and plotted in GraphPad Prism 9.1.0. Data is reported until the voltage of the buffer sample reached 400 V.

### Peptide crystallization

IQN17 was crystallized at room temperature in a hanging-drop vapor diffusion system by mixing 0.3 μL of 15 mg/mL peptide with 0.3 μL of reservoir solution containing 0.1 M imidazole (pH 8.0) and 35 % 2-methyl-2,4-pentanediol (MPD) (Qiagen). Crystals were seen two days after trays were set up. Single crystals were looped and directly frozen in liquid nitrogen. IQN22 was crystallized at room temperature in a hanging-drop vapor diffusion system by mixing 0.3 μL of 25 mg/mL peptide with 0.3 μL of reservoir solution containing 0.2 M di-ammonium phosphate and 40 % MPD. Crystals were observed one day after setting up trays. Single crystals were looped and directly frozen in liquid nitrogen.

### X-ray crystallography

For the IQN17 crystal, X-ray diffraction data was collected to 1.2 Å at the SLAC National Accelerator Laboratory Stanford Synchrotron Radiation Lightsource (SSRL) beam line 12-2. The crystal belonged to the space group *I* 2 2 2 with unit cell dimensions a = 45.8 Å, b = 46.9 Å, c = 136.2 Å, ⍺ = 90°, β = 90°, Y = 90°. For the IQN22 crystal, X-ray diffraction data was collected to 1.9 Å at SSRL beam line 14-1. The crystal belonged to the space group *P* 21 21 21 with unit cell dimensions a = 27.3 Å, b = 38.2 Å, c = 151.5 Å, ⍺ = 90°, β = 90°, Y = 90°.

Diffraction data for IQN17 were processed using HKL-3000.^62^ Diffraction data for IQN22 were processed using autoPROC^63^ to obtain the scaled and merged reflections. The structure of IQN17 was solved using Phaser in Phenix^64^ by molecular replacement with the low-resolution structure of IQN17 (PDB ID 2Q7C) as a search model. The structure of IQN22 was solved by molecular replacement with the apo IQN17 structure. Model improvements were achieved through iterative cycles of automated refinement in Phenix Refine^65^ combined with manual model fitting using Coot^66^ and validation using MolProbity.^67^ Structural images were generated with PyMOL (Schrödinger). Refinement parameters are summarized in Table S1.

### Oxidation of cysteine-containing peptides to form covalent trimers

Oxidation of peptides containing two cysteines at the N-terminus was adapted from previously described protocols.^28^ Briefly, lyophilized pure peptide was dissolved to approximately 0.7 mg/mL in 50 mM Tris-HCl (pH 8.0) and oxidized at room temperature with gentle shaking for 48-72 h. The solution was then lyophilized and re-purified by HPLC as described above. Product whose mass corresponded to three peptide chains linked by three disulfide bridges were collected. Side products corresponding to peptide dimers or peptide chains linked together by four disulfide bridges were observed.

### Mouse Immunizations

All mouse experiments were conducted at Josman LLC (Napa Valley, CA). Female Balb/c mice (6-8 weeks old) were immunized with 15 μg of peptide adjuvanted with 20 μg CpG (ODN1826, InvivoGen) and 500 μg Alhydrogel (InvivoGen) diluted in Dulbecco’s phosphate-buffered saline (DPBS) (Gibco). Mice were immunized on day 0 and boosted at day 21 and day 42. Serum was collected at days 0, day 21, day 35, day 63, and a final bleed was collected day 83.

### Enzyme-linked immunosorbent assay (ELISA)

MaxiSorp 96-well plates (ThermoFisher) were coated with antigen at 1 μg/mL in 50 mM bicarbonate pH 8.75 for 1 h at room temperature then blocked overnight at 4 °C with ChonBlock Blocking/Dilution ELISA buffer (Chondrex). In the case of biotinylated antigens, 96-well plates were first coated with streptavidin (Thermo Scientific) at 5 μg/mL in 50 mM bicarbonate pH 8.75 for 1 h at RT before blocking overnight with ChonBlock. ChonBlock was then manually removed before washing 3X with PBST (10 mM phosphate, 2.7 mM KCl, 0.05% Tween20, pH 7.4). Mouse serum samples were serially diluted in PBST with a starting concentration of 1:50 and then added to coated plates and incubated at room temperature for 1 h. Plates were washed 3X with PBST before adding HRP goat anti-mouse antibody (BioLegend 405306) at a 1:10,000 dilution in PBST. After incubation for 1 h at room temperature, plates were washed 6X with PBST and then developed for 5 min using One-Step Turbo TMB substrate (Pierce). Reactions were quenched with 2 M sulfuric acid before determining absorbance at 450 nm using a Synergy BioHTX plate reader (BioTek). The background value from wells containing no serum was averaged and subtracted from all wells on the plate.

Background-subtracted values were imported into GraphPad Prism 9.1.0 and fitted with a three-parameter nonlinear regression to obtain EC_50_ value. For the competition ELISA assay, D5_AR (200 nM) was preincubated on the plate with antigen for one hour before addition of mouse antisera.

### Antibody-avidity ELISA

Antibody avidity was assessed using a sodium thiocyanate (NaSCN) displacement ELISA. MaxiSorp plates were coated with streptavidin for 1 h at room temperature, washed with 3X with PBST, blocked with ChonBlock at 4 °C overnight and washed 3X with PBST as described above. After incubating plates with serially diluted mouse serum samples for 1 h at room temperature the plates were washed 3X with PBST and then 2 M NaSCN was added to each well. Plates were incubated for 15 min at room temperature before washing 3X with PBST. Secondary antibody, development, quenching, read out and analysis in GraphPad Prism to obtain EC50 values were performed as described above.

### Env-pseudotyped lentivirus production

Env-pseudotyped lentivirus was produced in HEK293T cells via calcium phosphate transfection as previously described.^68^ The day before transfection, 6 x 10^6^ HEK293T cells were seeded in 10 cm plates in 10 mL D10 medium (Dulbecco’s Modified Eagle Medium [DMEM] and additives: 10 % fetal bovine serum [Gemini Bio-Products], 2 mM L-glutamate, 1 % penicillin/streptomycin, and 10 mM HEPES, pH 7.0). Cells were incubated overnight at 37 °C and 5 % CO2 without shaking. For transfection, 20 μg of pSG3ΔEnv and 10 μg of HIV-1 Env sequence of interest were added to filter-sterilized water to a final volume of 500 μL followed by the addition of 500 μL of 2X HEPES-buffered saline (pH 7.0). The backbone pSG3ΔEnv plasmid was obtained through the NIH AIDS Reagent Program, from Drs. John C. Kappes and Xiaoyun Wu (Cat# 11051). To form transfection complexes, the solution was gently agitated while 100 μL of cold 2.5 M CaCl2 was added dropwise. The solution was incubated at room temperature for 20 min and then added dropwise to the plated cells. Approximately 12-18 h after transfection, the medium was removed and replaced with fresh D10. Virus-containing culture supernatants were harvested on day 3 post-transfection. Supernatants were centrifuged at 300 *g* for 5 min and filtered through a 0.45 μm polyvinylidene difluoride filter before aliquoting and storing at −80 °C.

### Viral neutralization assays

The neutralization assay was adapted from the TZM-bl assay for standard assessment of neutralizing antibodies against HIV-1 as previously described^43^. TZM-bl/FcγRI cells (HeLa luciferase/β-galactosidase reporter cell line stably expressing human CD4, CCR5, CXCR4, and FcγRI) obtained from the NIH AIDS Research and Reference Reagent Program as contributed by Drs. John C. Kappes and Xiaoyun Wu, were plated in white-walled, 96-well plates (Greiner Bio-One 655098) at a density of 5000 cells/well and incubated overnight at 37 °C with 5 % CO_2_. Mouse serum was heat-inactivated at 56 °C for 1 h and centrifuged at 3000 *g* for 10 min before dilution in D10 medium. HIV-1 Env-pseudotyped lentivirus in growth medium supplemented with DEAE-dextran (Millipore Sigma) was added to the serially diluted serum samples.Medium was removed from cells and replaced with 100 μL medium containing the diluted mouse serum and virus. After incubation at 37 °C for 48 h, BriteLite assay readout solution (PerkinElmer) was added to the cells and luminescence values were measured using a microtiter plate luminometer (BioTek) after shaking for 10 s. Neutralization titer was defined as the sample dilution at which the relative luminescence units (RLUs) were decreased by 50 % as compared to the RLUs of virus-only control wells after subtraction of background RLUs in wells containing cells only. Normalized values were fitted with a three-parameter nonlinear regression inhibitor curve in GraphPad Prism 9.1.0 to obtain IC_50_ value. Fits for serum neutralization assays were constrained to have a value of 0 % at the bottom. Neutralization assays were performed in biological duplicates with technical duplicates.

### Statistical Analysis

All normalization, curve-fitting, and statistical analyses were performed using GraphPad Prism 9.3.1 software. ELISA EC_50_ and neutralizing serum NT_50_ values were compared across groups at each time point using a Kruskal-Wallis ANOVA followed by a Dunn’s multiple comparisons test. *P* values less than 0.05 are considered significant and plotted. For all graphs * =*p* ≤ 0.05, ** =*p* ≤ 0.01.

## Data Availability

Coordinates and structure factors have been deposited in the RCSB Protein Data Bank (http://www.rcsb.org) under PBD ID codes 8F3A and 8F3B.

## Supplementary Information

This article contains supplementary information (Fig. S1-S7 and Table S1-S3).

## Acknowledgments

We thank all members of the Kim lab for helpful discussion and guidance during this project, especially MVF Interrante and Drs. AE Powell, PA Weidenbacher, M Sanyal, and BN Bell. We thank E Munro for assistance with data analysis. We thank Dr. J. S. Fraser for insightful discussion and technical expertise and staff scientists of the Stanford Synchrotron Radiation Lightsource (SSRL) especially Dr. T Doukov for support during X-ray crystallographic data collection. Use of the SSRL, SLAC National Accelerator Laboratory is supported by the US Department of Energy (DOE), Office of Science, Office of Basic Energy Sciences under Contract DE-AC02-76SF00515.

## Author contributions

TUJB, GE, LD, and PSK conceived the experiments. TUJB, ST, GE, and DF performed crystallization, structure determination and refinement. TUJB performed all other experiments. TUJB and PSK wrote the paper. All authors contributed to revising the manuscript.

## Supplemental Information

**Figure S1.**
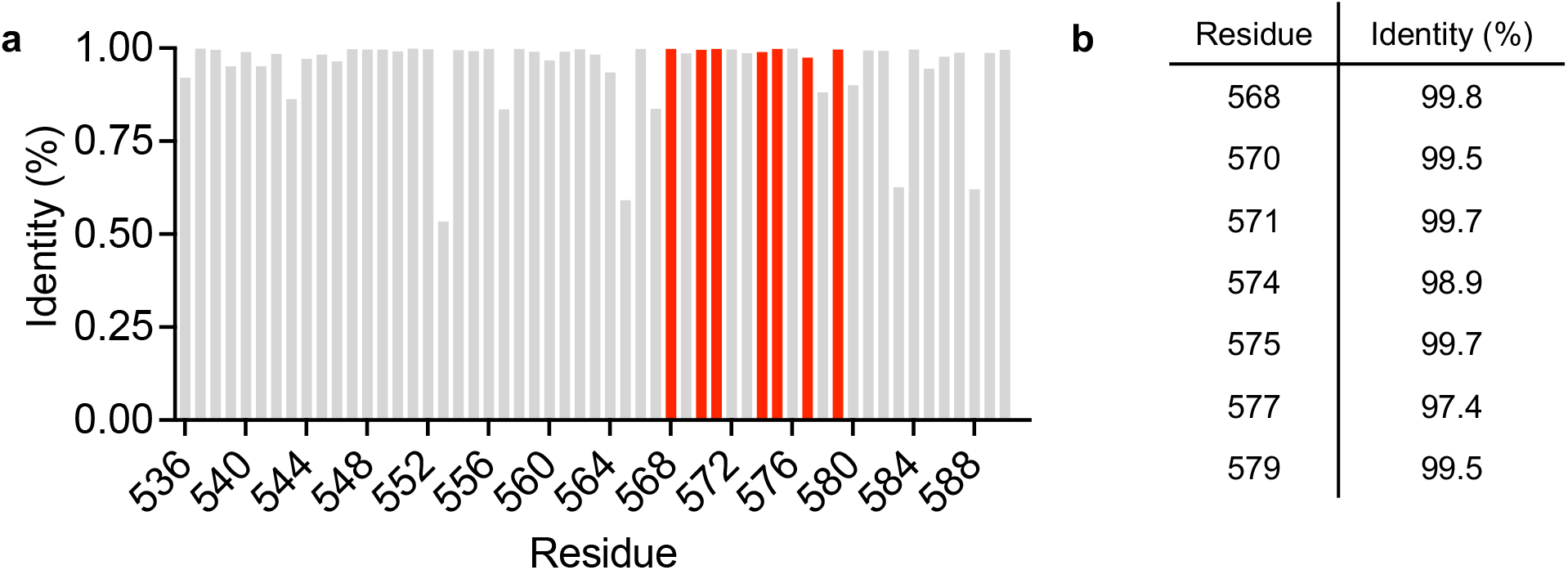
The seven residues that form the gp41 HIV-1 pocket are highly conserved across HIV-1 sequences. **a**, Percent identity at each position across the NHR of gp41 (HXB2 numbering) was calculated from the alignment of all complete HIV-1 sequences available in the Los Alamos database (https://www.hiv.lanl.gov/content/sequence/HIV/mainpage.html) as of October 2022. The seven residues forming the gp41 pocket are highlighted in red. **b,**Identity of the seven residues forming the conserved gp41 hydrophobic pocket.

**Figure S2.**
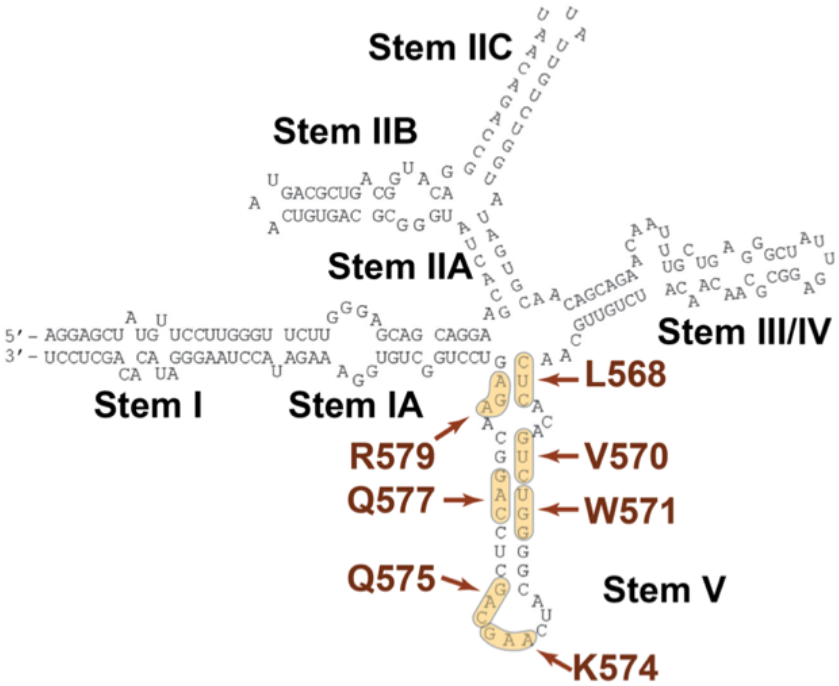
The mRNA nucleotides that encode the gp41 pocket residues are located in stem V of the HIV-1 rev response element. The seven highly conserved gp41 pocket residues are highlighted in brown (L568, V570, W571, K574, Q575, Q577, R579; HXB2 numbering).

**Figure S3.**
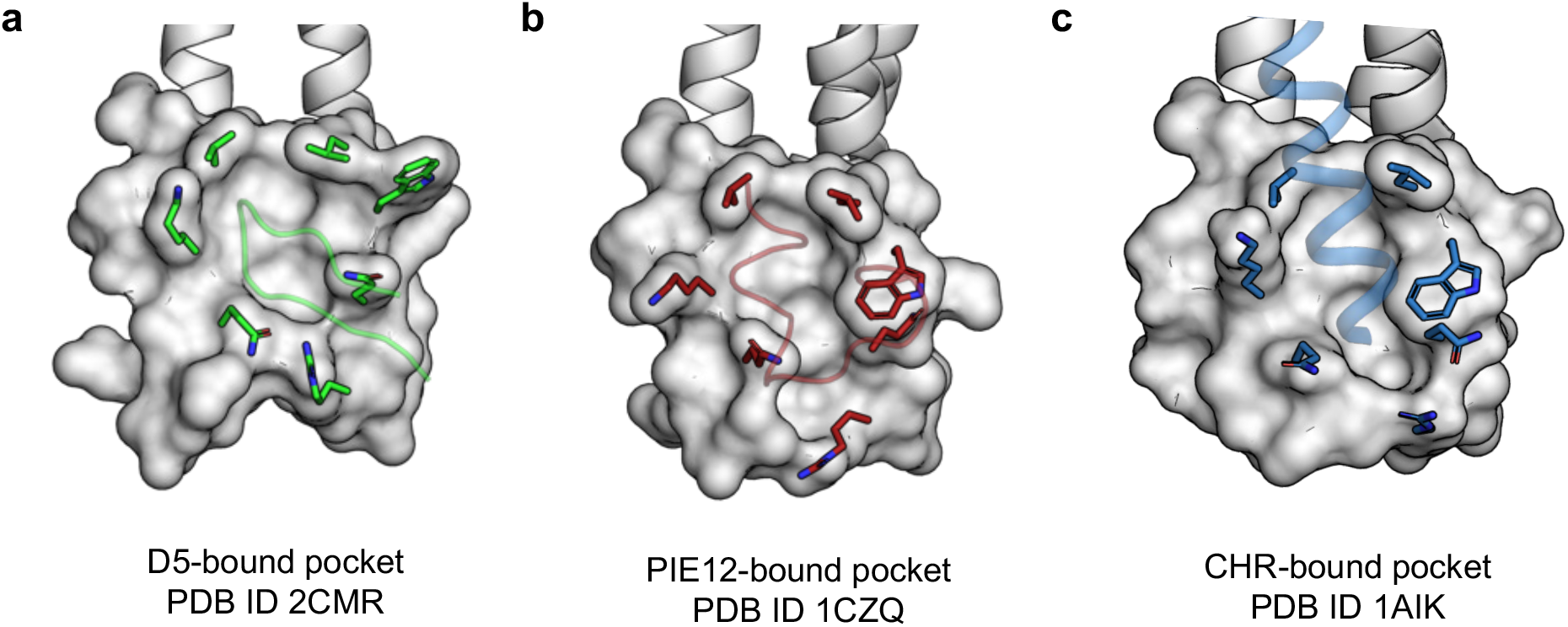
The gp41 pocket is highly malleable. Close up view of space-filling models of the gp41 pocket bound to a neutralizing antibody (green) (**a**), a D-peptide inhibitor (maroon) (**b**), or the CHR in the gp41 core structure (blue) (**c**). Pocket residues are shown as sticks.

**Figure S4.**
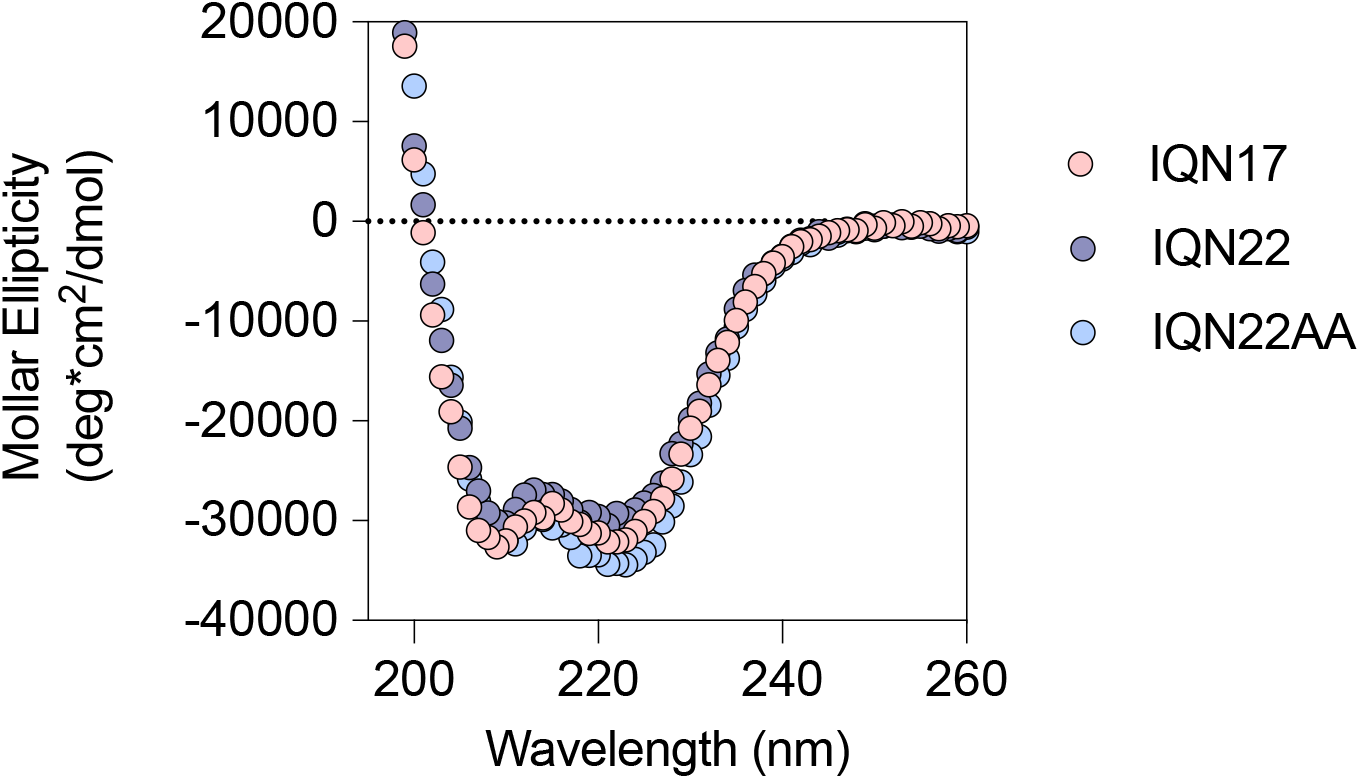
Circular dichroism (CD) spectra of the three covalently trimerized peptide immunogens. CD spectroscopy was obtained for HPLC-purified CCGG-containing peptides dissolved in PBS at a concentration of 20 μM. Each trace represents the average from three accumulations.

**Figure S5.**
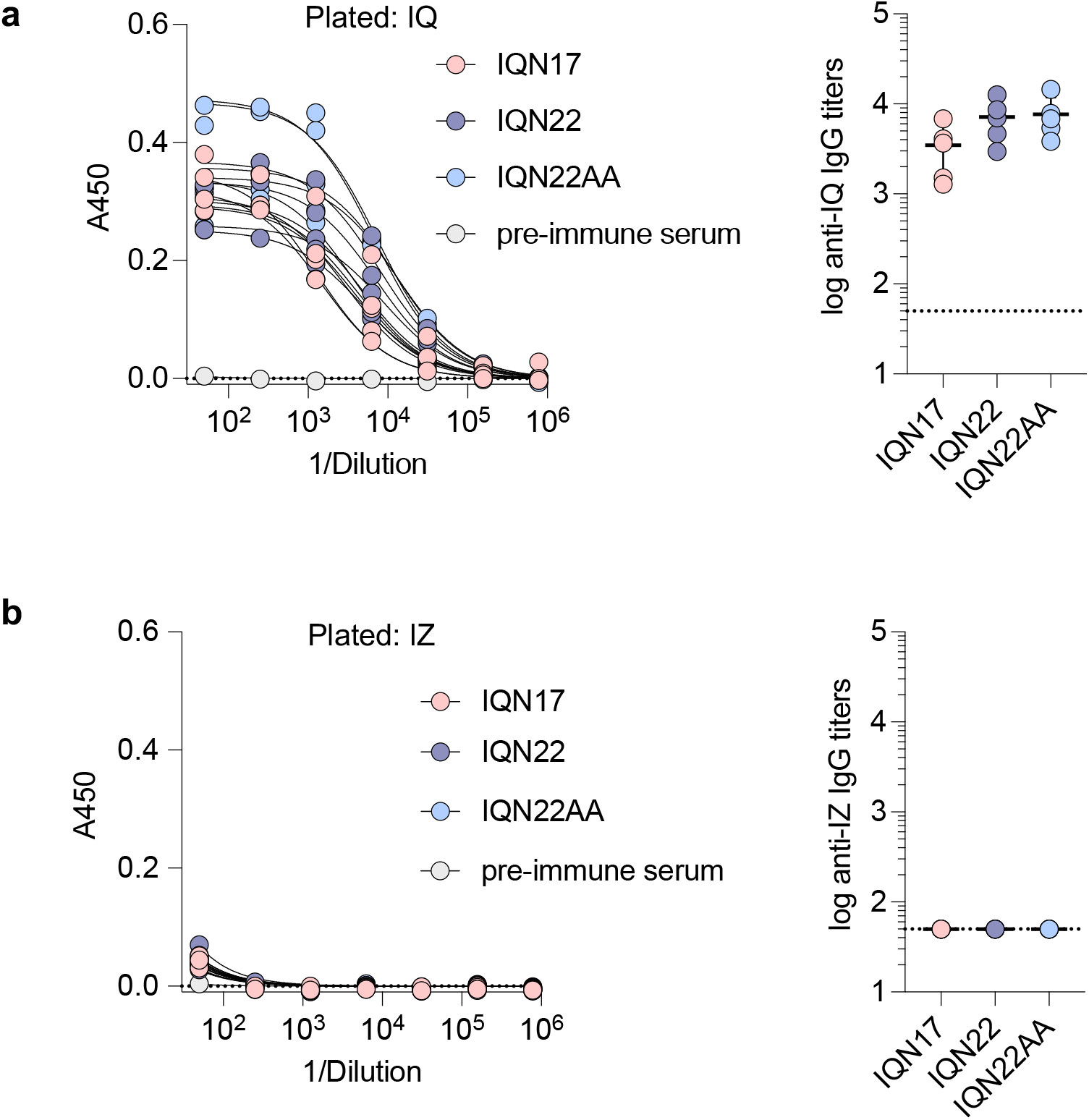
Immunized mice develop a response to the IQ trimerization domain but not the trimeric IZ domain. ELISA binding curves and binding titers of day 21 antisera from mice immunized with IQN17, IQN22, and IQN22AA (*n* = 5 per group) against the IQ domain (**a**) and the IZ domain (**b**). Serum binding titers are plotted as geometric mean ± s.d. of the log-transformed values. The horizontal dotted line represents the limit of quantitation.

**Figure S6.**
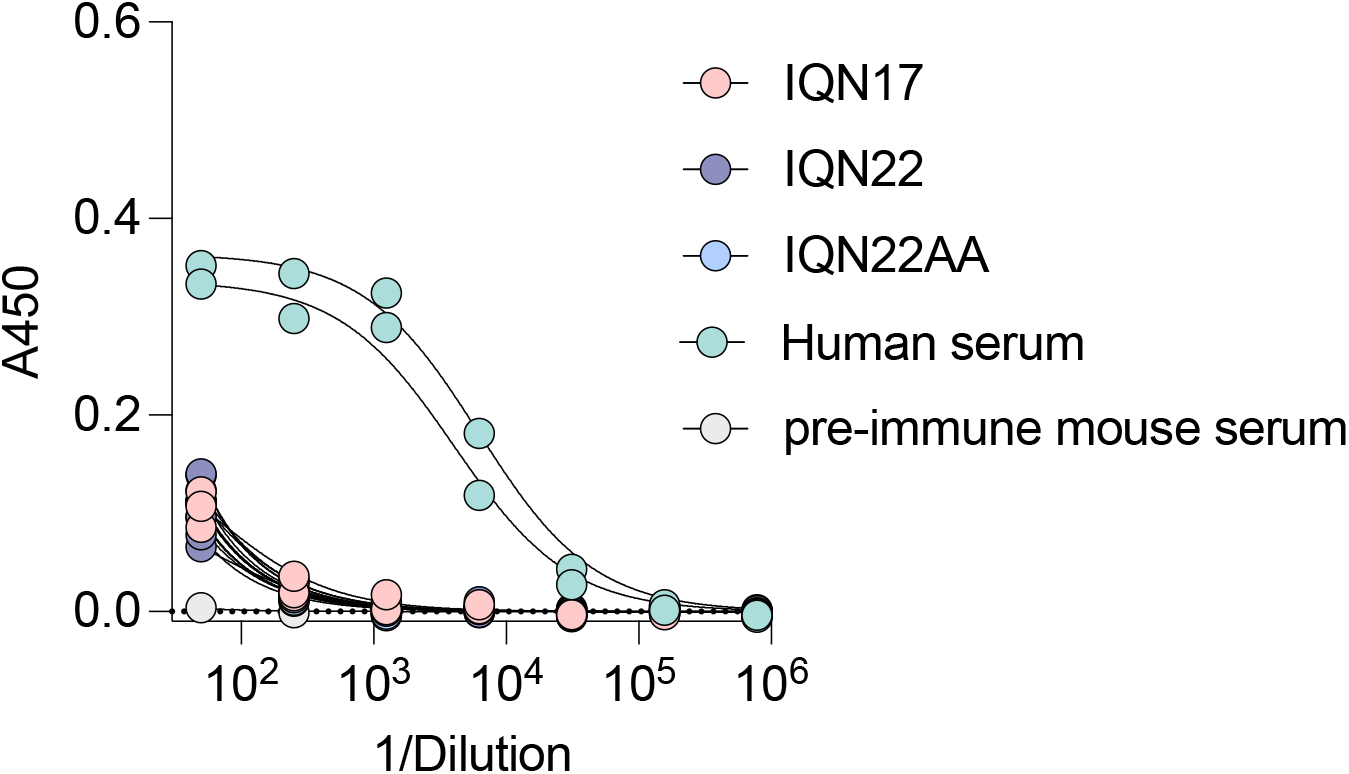
Immunized mice do not develop a response to the AVERY sequence of gp41. ELISA binding curves from mice immunized with IQN17, IQN22, and IQN22AA (*n =* 5 per group) against the peptide bio-AVERY. Human serum from HIV-1 positive individuals (*n* = 2) was used as a positive control.

**Figure S7.**
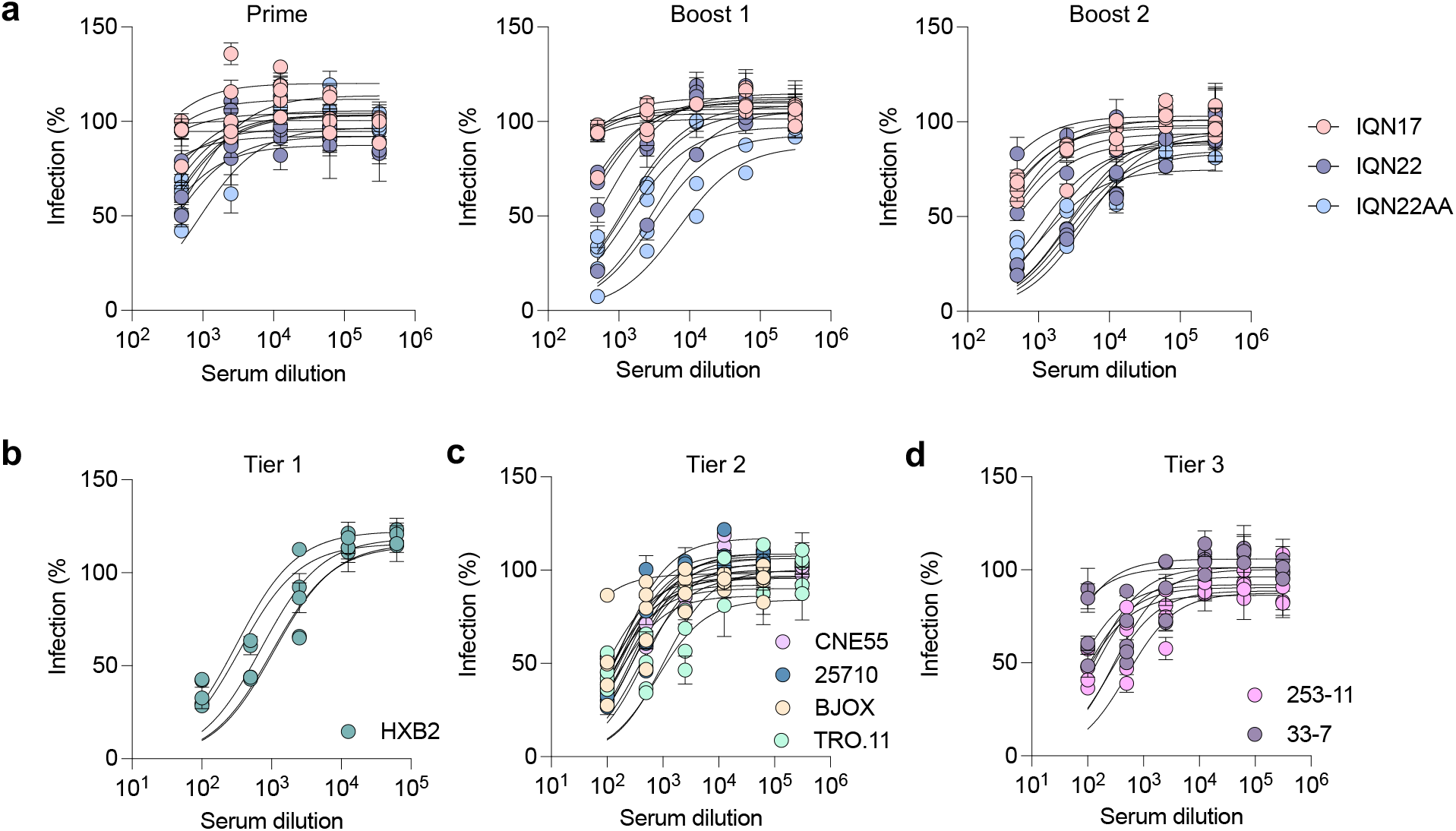
Neutralization of HIV-1 pseudoviruses. **a,**Serum neutralization of HXB2_L565Q_ by mice immunized with IQN17, IQN22, or IQN22AA at each time point. Serum neutralization of tier 1 (**b**), tier 2 (**c**), and tier 3 (**d**) viruses from mice immunized with IQN22AA post second boost. Neutralization assays were repeated in biological duplicate or triplicate. Representative neutralization curves from one assay are plotted as mean ± s.d. of technical duplicate.

**Table S1.** Crystallographic data collection and refinement statistics.

|  | IQN17 | IQN22 |
| --- | --- | --- |
| PDB ID code | 8F3A | 8F3B |
| Wavelength, Å | 0.979 | 0.979 |
| Resolution range, Å | 16.44 – 1.20 (1.24-1.20) | 37.05 – 1.98 (2.05-1.98) |
| Space group | *I* 2 2 2 | *P* 2<sub>1</sub> 2<sub>1</sub> 2<sub>1</sub> |
| a, b, c (Å) | 45.8 46.9 136.2 | 27.3 38.2 151.5 |
| α, β, γ (°) | 90 90 90 | 90 90 90 |
| Total reflections | 580,679 (55,278) | 151,616 (8,122) |
| Unique reflections | 46,268 (4,269) | 10,690 (1,108) |
| Multiplicity | 12.6 (12.2) | 14.1 (14.7) |
| Completeness, % | 98.83 (92.91) | 90.77 (98.04) |
| Mean I/sigma(I) | 25.1 (5.19) | 26.4 (2.3) |
| Wilson B-factor | 12.95 | 36.07 |
| *R*<sub>merge</sub> | 0.046 (0.381) | 0.064 (1.493) |
| *CC*<sub>1/2</sub> | 0.999 (0.993) | 1.00 (0.864) |
| *R*<sub>work</sub> | 0.17 (0.205) | 0.252 (0.322) |
| *R*<sub>free</sub> | 0.186 (0.247) | 0.259 (0.455) |
| No. of nonhydrogen atoms | 1425 | 1328 |
| Macromolecules | 1161 | 1240 |
| Water | 218 | 88 |
| Protein residues | 132 | 153 |
| RMS (bonds), Å | 0.019 | 0.019 |
| RMS (angles), ° | 1.42 | 1.25 |
| Ramachandran favored, % | 100 | 100 |
| Ramachandran outliers, % | 0 | 0 |
| Clashscore | 5.22 | 8.19 |
| Average B-factor | 20.15 | 49.16 |
| Macromolecules | 17.8 | 48.77 |
| Solvent | 32.7 | 54.6 |

Statistics for the highest resolution shell are shown in parenthesis.

**Table S2.** Average B-factors for all amino acids compared to R579 in IQN17 and IQN22.

|  | IQN17 |  | IQN22 |  |
| --- | --- | --- | --- | --- |
|  | All amino acids | R579 | All amino acids | R579 |
| **Cα** | 14 ± 3.5 | 21 ± 1.0 | 45 ± 10 | 45 ± 4.5 |
| **Backbone** | 14 ± 3.5 | 21 ± 1.3 | 44 ± 10 | 44 ± 5.0 |
| **Side chains** | 21 ± 7.0 | 36 ± 8.0 | 53 ± 13 | 48 ± 4.5 |

**Table S3.** Sequences of peptides used in this study.

| **Name** | **Sequence** |
| --- | --- |
| IQN17 | RMKQIEDKIEEIESKQKKIENEIARIKKLLQLTVWGIKQLQARIL |
| IQN22 | RMKQIEDKIEEIESKQKKIENEIARIKKLLQLTVWGIKQLQARILAVERY |
| IQN22AA | RMKQIEDKIEEIESKQKKIENEIARIKKLLQLTVWGIKQLQARILAVEAA |
| CCIQN17 | CCGGRMKQIEDKIEEIESKQKKIENEIARIKKLLQLTVWGIKQLQARIL |
| CCIQN22 | CCGGRMKQIEDKIEEIESKQKKIENEIARIKKLLQLTVWGIKQLQARILAVERY |
| CCIQN22AA | CCGGRMKQIEDKIEEIESKQKKIENEIARIKKLLQLTVWGIKQLQARILAVEAA |
| Bio-IQN17 | B*CCGGRMKQIEDKIEEIESKQKKIENEIARIKKLLQLTVWGIKQLQARIL |
| Bio-IQN22 | B*CCGGRMKQIEDKIEEIESKQKKIENEIARIKKLLQLTVWGIKQLQARILAVERY |
| Bio-IZN17 | B*CCGGIKKEIEAIKKEQEAIKKKIEAIEKLLQLTVWGIKQLQARIL |
| Bio-IZN22 | B*CCGGIKKEIEAIKKEQEAIKKKIEAIEKLLQLTVWGIKQLQARILAVERY |
| IQ | RMKQIEDKIEEIESKQKKIENEIARIKK |
| IZ | IKKEIEAIKKEQEAIKKKIEAIEKE |
| Bio-AVERY<sup>1</sup> | Bio[PEG]RILAVERYLKDQQLLGIWG |

B\* refers to the addition of biotin at the N-terminus by direct coupling to Biotin-PEG4-propionic acid (ChemPep).
<sup>1</sup>Peptide was purchased as part of a panel of gp41 peptides from Sigma-Aldrich.

